# Enhancing Plant Photosynthesis using Carbon Dots as Light Converter and Photosensitizer

**DOI:** 10.1101/2024.02.06.579025

**Authors:** Haitao Hu, Wenbo Cheng, Xueyun Wang, Yu Yang, Xuemeng Yu, Jianwei Ding, Yiliang Lin, Wei Zhao, Qiao Zhao, Rodrigo Ledesma-Amaro, Xihan Chen, Junzhong Liu, Chen Yang, Xiang Gao

**Author notes:** These authors contributed equally to this work (Haitao Hu, Wenbo Cheng, Xueyun Wang, Yu Yang, Xuemeng Yu).

## Abstract

Improving photosynthetic efficiency is pivotal for CO_2_-based biomanufacturing and agriculture purposes. Despite the progress on photosynthetic biohybrids integrating biocatalysts with synthetic materials, nanomaterials with improved optical and photoelectrochemical properties are still needed to increase the energy-conversion efficiency. Here, we present a novel approach using carbon dots (CDs) as both intracellular photosensitizers and light converters for enhancing solar energy utilization in photosynthetic organisms. The CDs were produced from cyanobacterial biomass and used to convert a broad spectrum of solar irradiation to red light. We demonstrated that the nanosized CDs were incorporated into cyanobacterial cells and transferred light-excited electrons into the photosynthetic electron transfer chain. The biohybrids consisting of the CDs and *Synechococcus elongatus* exhibited increased growth rates, enhanced activities of both photosystems, and accelerated linear electron transport, compared with the cyanobacterial cells only. The supplementation of the CDs increased CO_2_-fixation rate and CO_2_-to-glycerol production by 2.4-fold and 2.2-fold, respectively. Furthermore, the CDs were shown to enhance photosynthesis and promote growth of *Arabidopsis thaliana*. The fresh weight of plant was increased 1.8-fold by CDs addition. These results reveal that simultaneous photosensitization and spectral modification could substantially improve the efficiency of natural photosynthesis. This study presents CDs as an attractive nanomaterial with great application potential in agriculture and solar-powered biomanufacturing.

## Introduction

Photosynthesis, a process used by plants, algae, and cyanobacteria to convert CO_2_ and water into sugars with the aid of solar energy, provides food and energy for nearly all life on Earth^1^. Improving photosynthesis is crucial to feeding a growing population, especially with the negative effects of climate change and declining arable land^2, 3^. Although photosynthetic organisms have evolved efficient light-harvesting systems with a high quantum efficiency, the overall photosynthetic efficiencies are still low (i.e., 0.2–1% for crop plants)^4, 5^. During natural photosynthesis, light is absorbed by chlorophyll molecules in photosystems. As the chlorophylls are primarily sensitive to visible light, photosystems can only intercept about 40% of the incident solar energy^5^. Light reflection and transmission due to weak absorption of green light cause a further loss of energy. Attempts to improve photosynthesis included genetic manipulation of light-harvesting antenna^6^, engineering ribulose 1,5-bisphosphate carboxylase/oxygenase (Rubisco) for higher activity and specificity^7^, enhancing the regenerative capacity of the carbon reduction cycle^4^, and rewiring photorespiration to avoid CO_2_ release^8^. However, these efforts face challenges: firstly, only a few photosynthetic organisms are tractable for complex genetic engineering^9^. Additionally, the chlorophyll-based light harvesting and charge separation are kinetically restricted^10^, and those light-harvesting systems are prone to photodamage^11^, thereby impeding the efficiency of photosynthesis.

Photosynthetic biohybrids integrating biocatalysts with organic or metallic-based materials offer an alternative solution for efficient solar energy utilization^12–14^. In such hybrid systems, biocatalysts can be photosensitized with semiconductors that harvest light energy stably and efficiently^13, 15–18^. The photoexcited electrons are fed into proteins or cells to drive CO_2_ reduction or chemical production^15, 19, 20^. Previous studies have shown that semiconductor biohybrids can not only improve natural photosynthesis but also allow the light-driven conversion of CO_2_ to chemicals by chemolithoautotrophic bacteria^21–25^. Some organic polymers are capable of absorbing the light spectra that chlorophylls do not capture (e.g., ultraviolet and infrared light), thereby offering a promising solution to optimizing spectral quality for plants and other photosynthetic organisms^26, 27^. However, despite these progresses, no materials are yet capable of providing both spectral modification and photosensitization for improving photosynthesis. Such materials would have the potential to be highly useful in agriculture and solar-powered biomanufacturing.

Carbon dots (CDs), a type of carbon-based nanomaterials, have great advantages in terms of biocompatibility and low cost over metallic-based materials, especially considering the potential applications in agriculture at scale^28–30^. Importantly, CDs possess strong light absorption and outstanding capability of spectral modifications, resistance to photobleaching, and easy surface functionalization^31–33^. Thus, CDs have been used in diverse applications, including biomedicine (bioimaging, nanomedicine), sensors, anticounterfeiting, and energy (light-emitting diodes, solar cells, supercapacitors)^34–37^. Recently, CDs have been used as photosensitizers to transfer electrons to isolated fumarate reductase or hydrogenase for solar-driven catalysis ^38, 39^. Although a few studies have shown CDs as light converters to promote plant and algae growth^40, 41^, the applications of CDs in enhancing photosynthesis are still limited.

In this study, we introduce ultra-small and biocompatible CDs as intracellular photosensitizers and spectral converters for photosynthetic organisms, encompassing both plant and cyanobacteria (Fig. 1a). The ability of CDs to tune emission wavelength allows the conversion of ultraviolet (UV), blue, and green light to favorable red light, improving light utilization efficiency. These CDs generate an effective light-activated electrical current intracellularly, providing an additional supply of extra-exogenous electrons for the photosynthetic electron transfer chain (PETC) and avoiding inefficient electron transfer across the membrane. This work presents the successful integration of CDs with photosynthetic organisms to create CDs–biohybrid systems that significantly boost photosynthesis. In particular, the CDs–cyanobacteria hybrids showed enhancement in both the light-dependent reaction and the carbon fixation rate, leading to a more effective transformation of solar energy into chemical energy. Additionally, when using *Arabidopsis thaliana* as a model organism, we demonstrate that the CDs– biohybrid system facilitates faster plant growth. Our results suggest that biohybrid systems with biocompatible carbon-based photosensitizers holds significant potential for enhancing photosynthetic efficiency in various organisms, which can contribute to applications in sustainable agriculture and food safety.

**Fig. 1:**
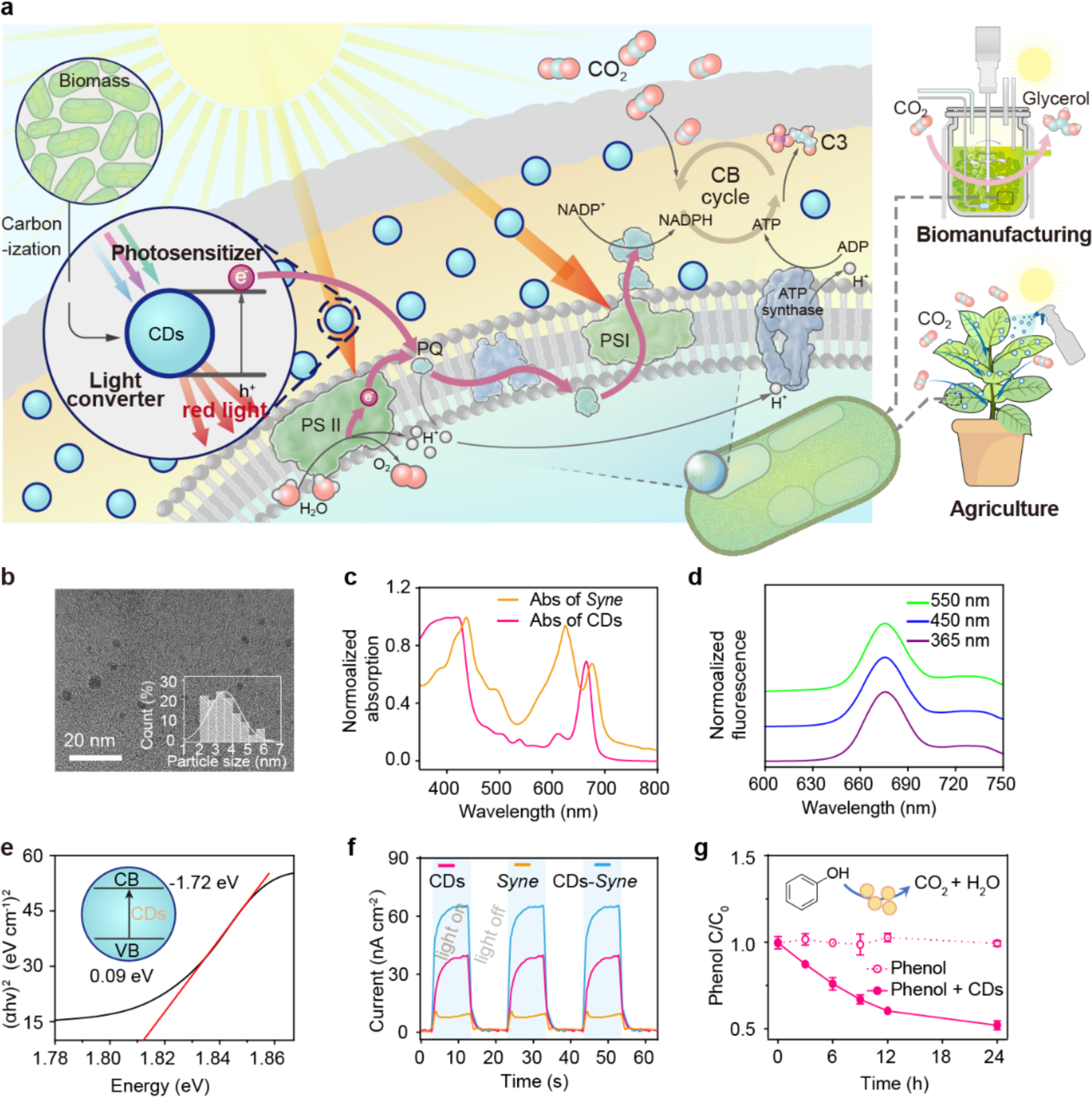
Schematic diagram of CDs for improving photosynthesis and characterization of CDs. **a.** Schematic of CDs for enhancing CO_2_-to-chemical production and plant biomass growth based on photosensitization and spectral modification. **b.** A representative TEM image of isolated CDs. The TEM images were used to analyze the diameter of particles (more than 110 particles). **c.** Normalized UV-visible absorption spectra of *Syne* and CDs. **d.** Normalized fluorescence spectra of CDs with UV, blue, and green light excitation. **e.** The Tauc plot for CDs. The intercept was used to calculate the direct bandgap. Inset: the band information of CDs. CB: conduction band, VB: valence band. α is the absorption coefficient, h is the Planck’s constant (6.626 × 10^−34^ J·s), and v is the frequency. **f.** A representative photocurrent curve of CDs, *Syne* cells, and CDs–*Syne* hybrids. **g.** The phenol degradation curve in the presence or absence of CDs under light condition.

## Results and Discussion

### Production and characterization of CDs

As plant or cyanobacterial biomass is one of the most abundant and low-cost feedstocks, we prepared the CDs using cyanobacterial biomass through hydrothermal method^42^ (Supplementary Fig. 1a). Cyanobacterial cells were collected and extracted with ethanol. After being heated in autoclaves, the solution was filtrated, dried, and extracted by dichloromethane. The CDs were thus obtained from the lower layer of the liquid. Transmission electron microscopy (TEM) analysis revealed the average size of the CD nanoparticles to be 3.5 nm (Fig. 1b). These particles mainly consisted of carbon (C), nitrogen (N), and oxygen (O) elements, and exhibited an amorphous structure (Supplementary Fig. 1b-f). The CDs exhibited a wide adsorption spectrum from 350 to 690 nm (Fig. 1c). By contrast, cyanobacterial and plant cells had limited adsorption of UV and green light^43^ (Fig. 1c). The red fluorescence emitted by the CDs spanned from 640 to 710 nm (maximum at 676 nm) (Fig. 1d), which were located within the optimal adsorption region of cyanobacteria and plants^44^. The emission λ_max_ remained nearly constant with excitation wavelengths at 365 nm, 450 nm and 550 nm, respectively (Fig. 1d), which is likely due to the surface states or defects emission^45^. These results demonstrate that these CD particles were capable of optimizing spectral quality.

We then measured the photoelectrochemical performance of the CDs, which possessed a direct bandgap energy of ∼ 1.81 eV with conduction band (CB) of −1.72 eV and valence band (VB) of ∼ 0.09 eV (Fig. 1e). The photoinduced current was ∼ 41 nA cm^−2^ (Fig. 1f). To test whether the photoinduced electron-hole charges separation by the CDs can catalyze the degradation of organic compounds, the intrinsic property of semiconductor, the CD particles were incubated with 100 mg L^−1^ of phenol. As PETC has a reducing potential higher than −1.3 eV^46^, the CDs could potentially reduce components in PETC. We observed that phenol concentration decreased by about 50% under illumination within 24 hours (Fig. 1g). In contrast, degradation of phenol was negligible in the absence of CD particles. Thus, CDs could potentially provide the exogenous electrons to the PETC and power the photosynthesis.

### Incorporation of CDs into *Synechococcus elongatus* cells

Due to the extremely small size of the CD particles (∼3.5 nm), we hypothesized that they could cross the cell membrane to enter the cell. Within the cell, the CD particles could generate photoexcited electrons, thus the energy-consuming transmembrane electron transfer could be bypassed^47^. To test our hypothesis, we added CDs to the cell suspension of a unicellular cyanobacterium *Synechococcus elongatus* UTEX 2973 (*Syne*). After incubation for 1 hour, the cells were collected and cross-sectional samples were prepared for high-angle annular dark field scanning transmission electron microscopy (HAADF-STEM) analysis. To enhance contrast for better imaging in HAADF-STEM, the CDs were doped with Zn (all CDs in other experiments did not contain Zn), which had no effect on the CDs’ morphology (Supplementary Fig. 2c). STEM-energy dispersive spectroscopy (EDS) mapping showed that the Zn elemental signal was in alignment with nitrogen and oxygen elements (biomass signals) and evenly distributed throughout the cells (bottom of Fig. 2a, and Supplementary Fig. 2a). In contrast, no Zn was detected for control samples without CDs (top of Fig. 2a). The STEM data indicated that the CDs were efficiently translocated into the *Syne* cells within 1 hour, resulting in the formation of CDs–*Syne* biohybrids. In addition, the zeta potentials of *Syne* shifted from −14 mV to −21 mV upon combination with CDs, which had a zeta potential close to −1 mV (Fig. 2b). Importantly, the CDs–*Syne* displayed a peak photocurrent of 62 nA cm^−2^, surpassing the sum of the photocurrents from CDs only (41 nA cm^−2^) and *Syne* cells only (9 nA cm^−2^) (Fig. 2b). These results indicated the successful incorporation of CDs into *Syne* cells.

**Fig. 2:**
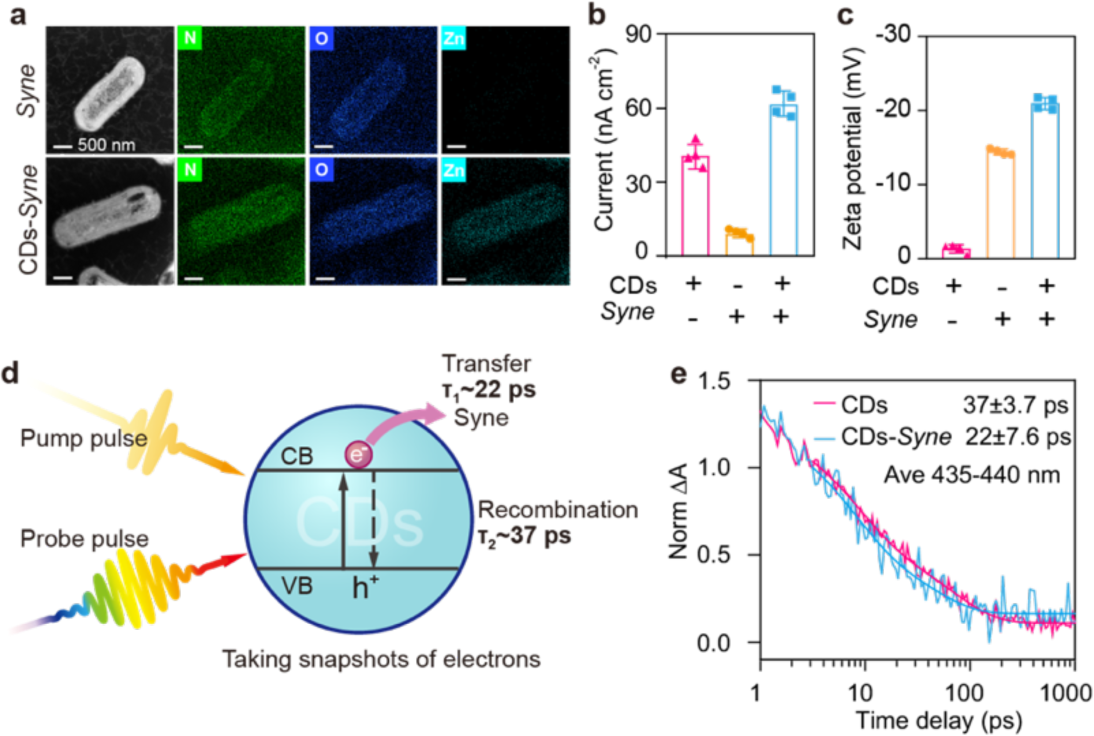
Characterization of the formed CDs–*Syne* hybrids. **a.** Cross-sectional STEM image of biohybrid sample. **b.** The photocurrent of CDs, *Syne* cells, and CDs-*Syne* hybrids. **c.** The zeta potentials of CDs, *Syne* cells, and CDs–*Syne* hybrids. **d.** Schematic of transient absorption (TA) spectroscopy analysis. τ_1_ and τ_2_ represent the charge transfer process and charge recombination process, respectively. **e.** Transient kinetics for pure CDs and CDs-*Syne* hybrids.

To study the charge transfer between CDs and *Syne* interfaces, we performed transient absorption (TA) spectroscopy analysis. The CDs were excited by the pump light and the electron dynamics signals were recorded by the probe light (Fig. 2d). We firstly explored whether the observed dynamics signals were electron dominates. TA spectra of CDs with electron capture agent (Fe^3+^) or hole trapping agent (Vitamin c) were shown in Supplementary Fig. 3a and b. The TA spectra of CDs (430 nm and 450 nm) were almost overlapped with the vitamin addition samples, but were different from the Fe^3+^ addition samples. Those results confirmed the dynamic signals in the 435–440 nm range, in which the electron signal dominates. The TA spectrum of CDs–*Syne* hybrids exhibited a faster decay than CDs only (Fig. 2e). This result suggests that the presence of *Syne* facilitates electron transfer from CDs. To quantitatively analyse the charge transfer rate, a biexponential function was employed^48, 49^. The charge transfer process in the CDs–*Syne* took only 22 ps, which is approximately a 40% decrease compared to that of the CDs only (37 ps). Overall, these results demonstrate that photoexcited electrons were transferred from CDs to the *Syne* cells at a rate of a few ps.

### Enhanced photosystem activity and carbon fixation in CDs–*Syne* hybrids

To assess whether the CDs could improve the efficiency of photosynthesis, we cultivated *Syne* cells in minimal medium with varying concentrations of CDs (10, 40, 80, and 160 mg L^−1^). Cell growth was monitored by measuring optical density at 730 nm (OD_730_). The CDs–*Syne* hybrids, compared with the *Syne* cells only, showed accelerated growth with CD concentrations lower than 80 mg L^−1^ (Fig. 3a). The observed growth inhibition at a high CD concentration (160 mg L^−1^) (Fig. 3a) was likely due to the generation of reactive oxygen species (ROS)^50^. Notably, at the optimal concentration of CDs (40 mg L^−1^), the growth rate of *Syne* cells reached 0.1 h^−1^ during the exponential growth phase, which was 1.5-fold higher than that of the control group without CDs. As photoautotrophs, *Syne* cells fix and convert CO_2_ into biomass, and they derive energy from photosynthesis for CO_2_ fixation and biomass formation^51^. Thus, our results suggest that the CDs enhance the photosystem activity and CO_2_ fixation of *Syne* cells.

**Fig. 3:**
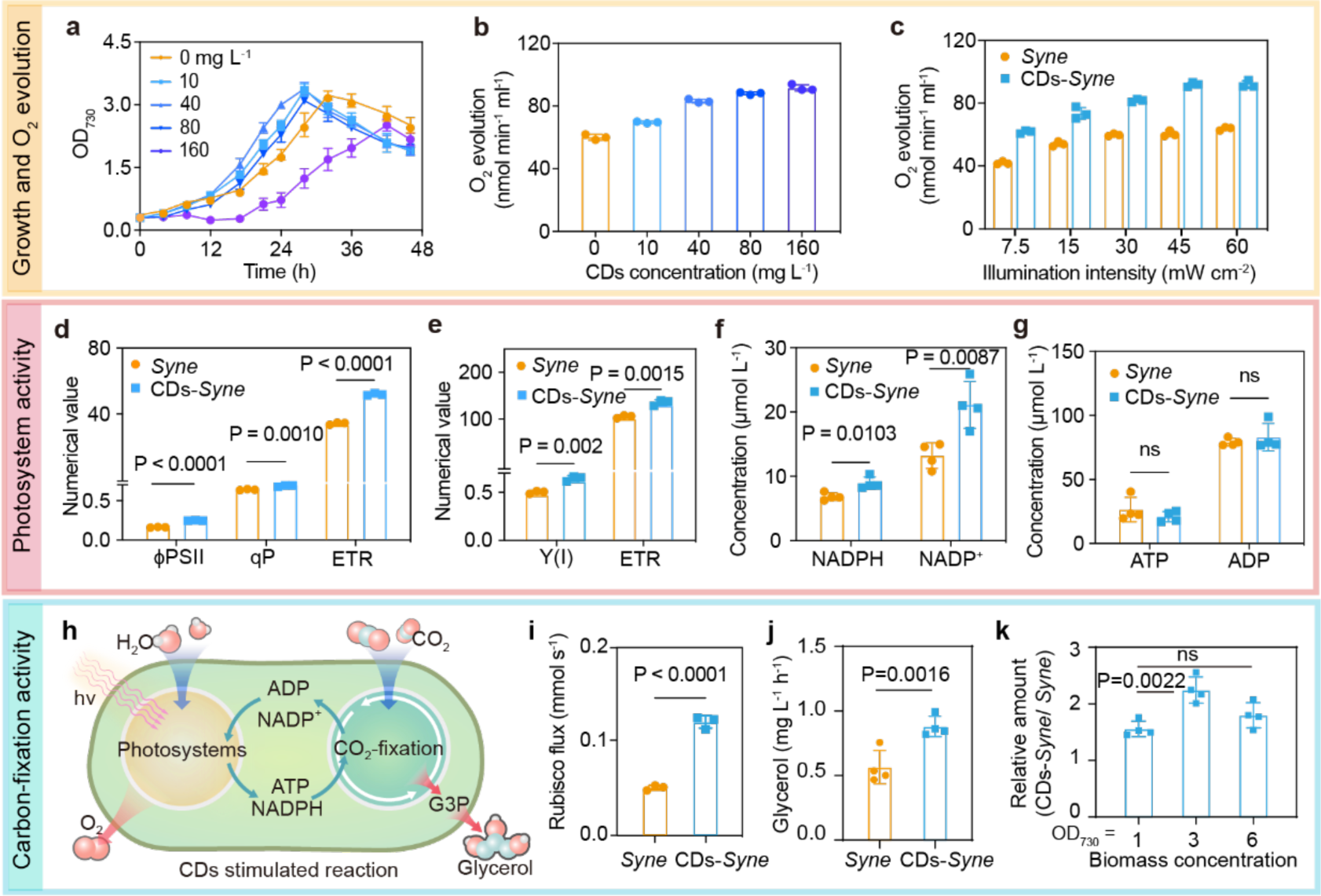
Enhancement of cyanobacterial photosynthesis by CDs. **a.** The growth curve of *Syne* cells with different concentrations (10, 40, 80, and 160 mg L^−1^) of CDs. **b.** The oxygen evolution rate of *Syne* cells with different concentrations of CDs under light intensity of 30 mW cm^−2^. **c.** The oxygen evolution rate of *Syne* cells with 40 mg L^−1^ CDs under various light intensity. **d.** Quantum yield (ϕPSII), photochemical quenching coefficient (qP) and electron transport rate (ETR) of PSII in CDs–*Syne* hybrids and *Syne* cells only. **e.** Quantum yield (Y (I)) and ETR of PSI in *Syne* cells and CDs–*Syne* hybrids. **f.** The intracellular concentration of NADP^+^ and NADPH in *Syne* cells and CDs– *Syne* hybrids. **g.** The intracellular concentration of ADP and ATP in *Syne* cells and CDs–*Syne* hybrids. **h.** Schematic of CDs-enhanced activities of photosystems for ATP and NADPH generation and CO_2_ fixation in cyanobacterial cells. **i.** The metabolic flux through Rubisco in *Syne* cells and CDs–*Syne* hybrids. **j.** Glycerol production in *Syne* cells and CDs–*Syne* hybrids under white light LED (typically red, green, and blue (RGB)) condition for 1 hour. **k.** Relative glycerol production by CDs–*Syne* hybrids compared to that by *Syne* cells under different cell densities (OD_730_=1, 3, and 6).

We then studied the effect of CD supplementation on the activities of cyanobacterial water-splitting oxygen-producing photosystems that generate reducing power NADPH and a proton gradient for the regeneration of ATP^52^. An increase in the photosynthetic oxygen evolution was observed upon CD addition, regardless of the light intensity conditions, low or high (Fig. 3b and c). The oxygen evolution rate was enhanced with increasing amounts of the CDs (Fig. 3b). In the presence of 160 mg L^−1^ CDs, the oxygen production rate reached 91.7 nmol min^−1^ mL^−1^, which was increased by 52.5% compared to the control without CDs (60.1 nm min^−1^ mL^−1^). Furthermore, the chlorophyll fluorescence and P700 of *Syne* were measured using a Dual-PAM-100 fluorometer to assess the activities of PSII and PSI. Compared with the control without CDs, the effective quantum yield of PSII (ϕPSII) in the CDs–*Syne* hybrids increased by 51.7% (Fig. 3d), which indicated an increase in the proportion of PSII-absorbed light energy that was used for charge separation and water oxidation. The photochemical quenching coefficient (qP) of *Syne*, which represents the proportion of open PSII reaction centers, was increased by 5% when incubated with the CDs (Fig. 3d). The CDs enhanced the electron transport rate (ETR) of PSII in *Syne* by 51.5%. Therefore, the CDs improved the PSII activity of *Syne*. Based on P700 measurements, we found that the effective quantum yield of PSI, Y(I), in the CDs–*Syne* hybrids was 31.1% higher than that in the *Syne* cells (Fig. 3e). The ETR of PSI in the *Syne* was increased 1.31-fold when incubated with CDs. Therefore, the CDs enhanced the activities of both photosystems and accelerated the linear electron transport in *Syne*. In addition, the intracellular NADPH concentrations in *Syne* were also measured, which were raised from 6.87 to 8.94 µM (30% increase) by incubation with CDs (Fig. 3f). The intracellular pool size of ATP was not changed significantly (Fig. 3g), which could be explained by a simultaneous increase in ATP production and consumption for biomass synthesis^53^.

We next examined the effect of CD supplementation on cyanobacterial carbon fixation activities. The metabolic flux through Rubisco in the Calvin-Benson (CB) cycle was determined based on [^13^C]-bicarbonate dynamic labelling experiments and measurements of the incorporation of ^13^C into the reaction product, 3-phosphoglyceric acid (PGA; Fig. 3h)^54^. The CO_2_-fixation flux was 0.146 mM s^−1^ in the CDs–*Syne* hybrids, which was 2.4-fold higher than that in *Syne* cells (0.06 mM s^−1^, Fig. 3i). Moreover, the intracellular PGA concentration in the CDs–*Syne* hybrids increased 2-fold (Supplementary Fig. 4a), and the concentrations of other CB cycle intermediates including glucose-6-phosphate and fructose-6-phosphate also increased when compared with the control without CDs (Supplementary Fig. 4b). These results suggest that the CDs augmented the photosynthetic CO_2_ fixation in *Syne* (Fig. 3h).

To investigate whether the CDs–biohybrids allow efficient photosynthetic production of chemicals from CO_2_, we added the CDs to a cell suspension (OD_730_ ∼ 1) of a glycerol-producing *S. elongatus* strain XG608^48^. After incubation for one hour under white light LED (1.35 mW cm^−2^), the CDs–*Syne* hybrids produced glycerol from CO_2_ at a rate of 0.57 mg L^−1^ h^−1^, which was 56% higher than that from XG608 without CDs (Fig. 3j). This improvement was notably higher than that reported for the Au–*Syne* hybrids^48^. In addition, we observed that CD supplementation induced a 2.2-fold increase in the glycerol production from *Syne* at a relatively high cell density (OD_730_ ∼ 3) (Fig. 3k). As the cell density increased, the light intensity for individual *Syne* cells decreased. Consistently, under a high light intensity (6.00 mW cm^−2^), the glycerol production from the CDs–*Syne* hybrids increased by only 18% compared to that from the *Syne* cells (Supplementary Fig. 5). We can therefore conclude that CDs improved the cyanobacterial chemical production from CO_2_, particularly under low light intensities.

### CDs enhance photosynthesis via photosensitization and light conversion

To determine whether the photoexcited electron transfer from the CDs contributed to the enhancement of cyanobacterial photosynthesis, we examined the effects of photosynthesis inhibitors on the glycerol production from the CDs–*Syne* hybrids. The inhibitors tested were 3-(3,4-dichlorophenyl)-1,1-dimethylurea (DCMU), 2,5-dibromo-3-methyl-6-isopropylbenzoquinone (DBMIB), and phenylmercuric acetate (PMA)^55^ (Fig. 4a). We found that in the presence of DCMU, which inhibits the electron transport from the PSII reaction center to plastoquinone (PQ), CDs supplementation still resulted in a 3.3-fold increase in the cyanobacterial glycerol production (Fig. 4b). By contrast, the glycerol production was not changed significantly by CDs in the presence DBMIB, which inhibits the electron flow from the PQ to cytochrome *b*_6_f, or PMA, which inhibits ferredoxin and NADP^+^ reduction (Fig. 4b). These results indicated that the CDs transferred photoexcited electrons to the PETC probably via PQ, which is consistent with previous studies that have shown the entry of exogenous electrons into the PETC through PQ^55^.

**Fig. 4:**
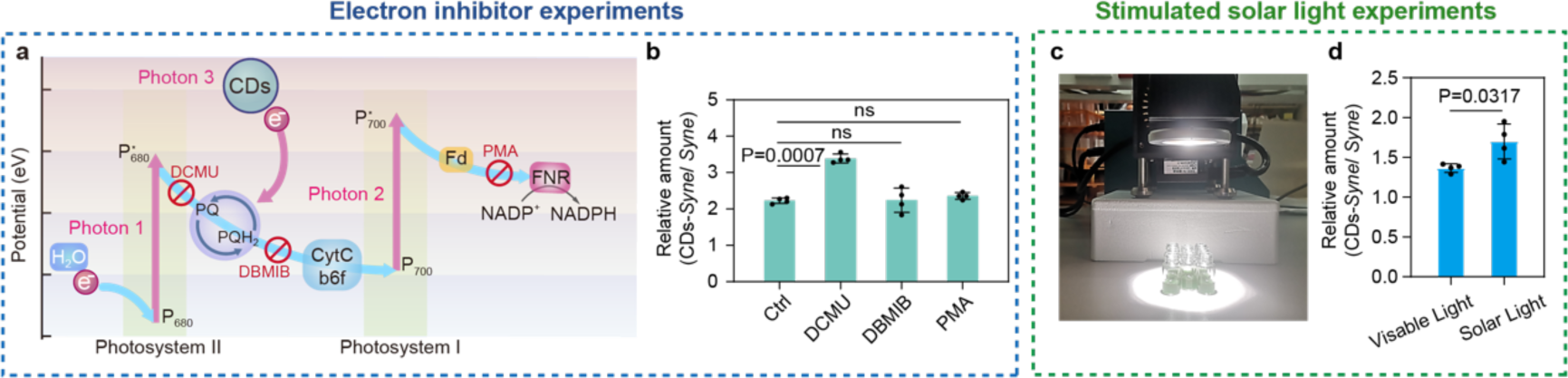
CDs enhance photosynthesis via photosensitization and light conversion. **a.** Schematic of photoexcited electron transfer from CDs to PETC. The crosses indicate the blocking site of individual photosynthesis inhibitors. Fd, ferredoxin; FNR, ferredoxin-NADP^+^ reductase. **b.** Relative glycerol production by CDs-*Syne* hybrids compared to that by *Syne* cells in the presence of photosynthesis inhibitors. **c.** Experimental set-up of simulated solar light irradiation by Xe lamp. **d.** Relative glycerol production by CDs-*Syne* hybrids compared to that by *Syne* cells under simulated solar light irradiation and visible light irradiation.

Given that CDs can behave as light converters, we studied the effect of their supplementation on cyanobacterial photosynthesis under simulated solar light irradiation using a Xe lamp (Fig. 4c). The CDs–*Syne* hybrids were incubated under the same intensity (1.5 mW cm^−2^) of either visible light or simulated solar light irradiation. A 1.7-fold increase in glycerol production was caused by the addition of CDs under stimulated solar light, which was 24% higher than that under visible light (Fig. 4d). This is probably due to the conversion of UV or green radiation to photosynthetically active radiation. Therefore, the spectral modification and photosensitization by the CDs enabled an enhancement of cyanobacterial photosynthetic efficiency.

### CDs improve photosynthetic efficiency in *A. thaliana*

To determine whether the CDs could also enhance plant photosynthesis, we tested the effect of CDs addition on the growth of *A. thaliana*. Various concentrations of CDs were sprayed on the leaves of *A. thaliana* seedlings which were cultivated under LED light. The fresh weight and leaf area were measured after two weeks of cultivation. We observed that the CDs could successfully accumulate in the leaves of *A. thaliana* (Fig. 5a). No growth inhibition of *A. thaliana* was observed at all CD concentrations tested, indicating a good biocompatibility of CDs. The supplementation of high concentrations of CDs (50 or 100 mg L^−1^) resulted in substantially improved plant growth, though the effect of a low concentration (10 mg L^−1^) of CDs was not significant (Fig. 5b). The plant growth was enhanced with increasing amounts of CDs. At 100 mg L^−1^ CDs, the fresh weight of *A. thaliana* reached about 0.42 g, which was over 1.8-fold higher than the control without CDs (Fig. 5c). The leaf area was increased from 9.1 cm^2^ to 12.7 cm^2^ by 100 mg L^−1^ CDs (Fig. 5d). These results indicate that the CDs enhanced photosynthesis and promoted growth of *A. thaliana*.

**Fig. 5:**
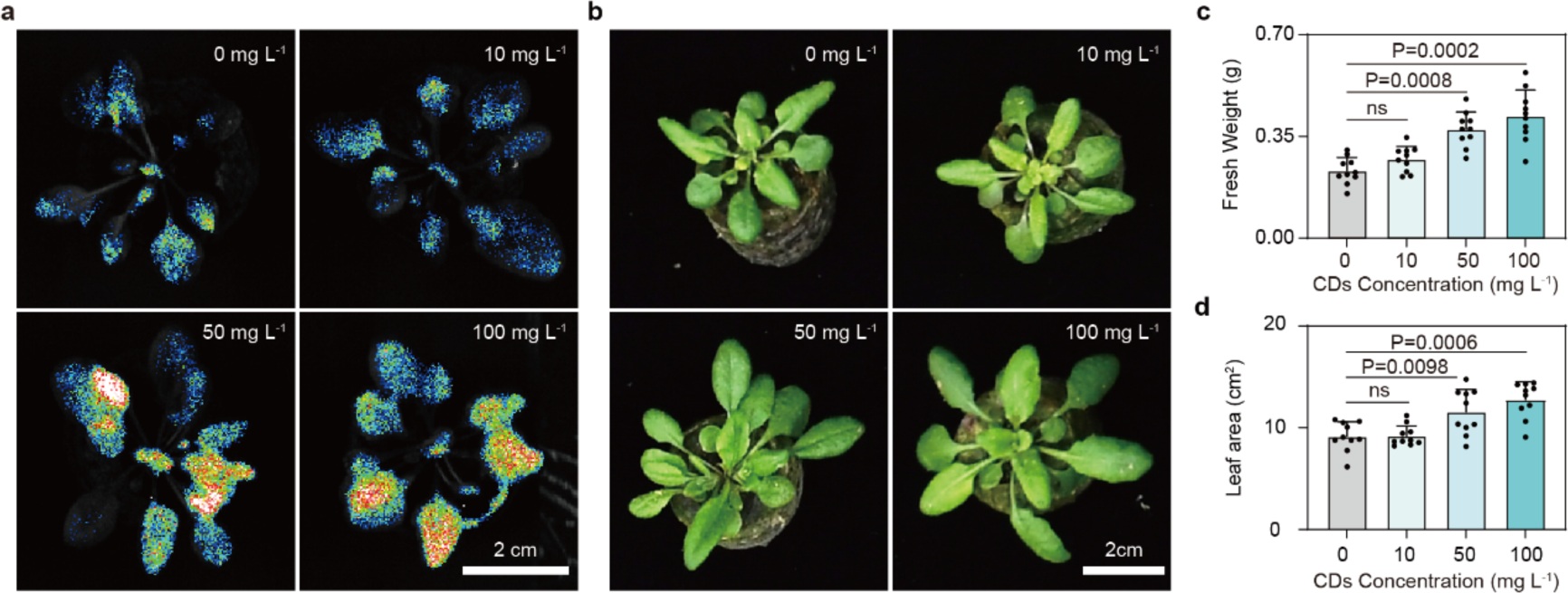
CDs enhance photosynthesis and promote growth of *A. thaliana*. **a.** The fluorescence photographs of *A. thaliana* growth with different concentrations of CDs at 7 days post the treatment. **b-d.** The photographs (b), fresh weight (c), and leaf area (d) of *A. thaliana* at 14 days post the treatment with different concentrations of CDs (n = 10).

## Conclusion

In summary, we have shown that CDs, acting as a light converter and intracellular photosensitizer, could increase photosynthesis efficiency in both cyanobacteria and higher plant. The biomass-based biocompatible CDs have a strong and wide absorption spectrum and a constant red-emission, which improve the quantity and quality of photons, promoting natural photosystems. Moreover, the photoinduced electrons from intracellular CDs could be passed to the PETC for producing additional reducing energy. In addition, the plant biomass production is increased by 1.8-fold and the solar-to-chemical efficiency by 2.2-fold in the CDs–biohybrid system. This work highlights the potential of CDs to provide a higher solar energy utilization and to increase crop yield and biochemical production.

## Methods

### Culture condition of *S. elongatus*

The shake flask cultures of *S. elongatus* PCC 2973, 7942 and derivatives were performed at 30℃ under continuous LED white light (1.35 mW cm^−2^) in 100 mL flasks with 30 mL of BG11 medium containing 100 mM NaHCO_3_ and 20 mM 2-[4-(2-hydroxyethyl)piperazin-1-yl]ethanesulfonic acid (HEPES).

### Preparation of CDs and CDs doped with Zn (Zn-doped-CDs)

*Syne* cells were cultured to an OD_730_ of 3 in BG11 medium. Cells pellets were collected and extracted in ethanol (1:10, w/v, 4 hours in dark). Afterwards, the cell debris was removed by filtration (0.22 μm filter paper), and the filtrate was used for synthesis the CDs through hydrothermal method^42^. Transferred the solution to a poly (tetrafluoroethylene) lined autoclave (50 mL) and heated in an oven at 150 ℃ for 4 hours. After cooling down to ambient temperature, the solution was filtrated through a 0.22 μm membrane and dried by a rotary evaporator. The crude product was extracted in a mixed solution of dichloromethane and water with a volume ratio of 1:1. After standing and layering, collected the dark green lower liquid layer and dried with nitrogen gas to obtain the CDs. CDs solid was diluted with ethanol to a suitable concentration for further use.

As for Zn-doped-CDs, 100 mM ZnCl_2_ was added to the filtrate before the hydrothermal reaction, and the following processes were the same as above.

### TEM of CDs

For the morphology characterization of CDs or Zn-doped-CDs, 1 mg mL^−1^ material in ethanol was dropped and dried onto a copper grid of 200 meshes, then TEM was detected on JEOL JEM F200 (Japan) at 200 kV.

### XRD of CDs

To determine the crystal structure of CDs, X-ray diffraction (XRD) was detected on Rigaku Ultima IV (Japan) at 10∼80°.

### XPS of CDs

To distinguish the elemental composition of CDs, X-ray photoelectron spectroscopy (XPS) was on Thermo Scientific K-Alpha (America) using Al Kα-ray at a work voltage of 12 kV using 1486.6 eV energy, and the data were calibrated using C1s at 284.80 eV.

### UPS and UV-vis of CDs

Ultraviolet photoelectron spectroscopy (UPS, ThermoFisher Nexsa) was used to detect the work function and valence band of CDs, and UV-vis (HORIBA Fluorolog^®^-3 Spectrofluorometer, America) was used to measure the direct bandgap of CDs and the absorption and fluorescence spectra of samples.

### Photocatalytic degradation of phenol using CDs

1000 mg L^−1^ CDs and 100 mg L^−1^ phenol were added into the 4 mL quartz reactors. The photocatalytic degradation reactions were performed at 25℃ under 12 mW cm^−2^ blue light LED. The samples were collected from quartz reactor, and the residue phenol concentrations were detected by ^1^H NMR (Bruker, 400 MHz). 100 mg L^−1^ phenol solution without CDs under the sample condition was used as the control.

### CDs–*Syne* hybrid construction for photocurrent and electron microscope image

*Syne* cells were inoculated with an initial OD_730_ of 0.2 in BG11 medium. IPTG (1 mM) was added to the cultures at OD_730_ of 0.4 if need. After cells were grown for 36 hours, the *Syne* cells were collected, washed, and resuspended in fresh BG11 medium to an OD_730_ about 1.5∼2.0. Added 100 mg L^−1^ CDs into the *Syne* cultures and incubated 1hour for photocurrent and electron microscope image analysis.

### CDs–*Syne* hybrid resin slice preparation and TEM characterization

*Syne* was cultured to the OD_730_ of 2, added 500 mg L^−1^ Zn-doped-CDs and incubated for 1 hour. The cell pellets were collected and washed with water twice, and then fixed with 2.5% glutaraldehyde overnight at 4℃. *Syne* without Zn-doped-CDs under the same conditions was used as the control.

Samples were washed with PBS (0.1 M, pH 7.0) 3 times for 15 min each, treated with 1% osmium tetroxide in PBS (0.1 M, pH 7.0) for 1-2 h and then rinsed in PBS (0.1 M, pH 7.0) three times for 15 min each. For dehydration, the samples were treated with a series of ethanol (30%, 50%, 70% and 80%) for 15 min, then exposed in acetone (90%, 95%) for 15 min and acetone (100%, 100%) for 20 min each. When preparing resin embedded samples, samples were exposed in acetone/Spurr agent (1,2,3-propanediol dehydrated glycerol ether) (1:1, v/v) for 1 h, acetone/Spurr agent (1:3) for 1 h and pure Spurr agent overnight. The samples were transferred into 1.5 mL EP tubes containing pure Spurr agent, the embedded resin blocks were obtained after 9 h at 70℃. The resin blocks were cut to 70∼90 nm ultrathin section by ultra-microtome (Leica UC7, German) with diamond knife (Daitome Ultra 45°), the sections were fished out onto copper grids of 200 meshes, then stained with both uranyl acetate for 8-15 minutes and lead citrate for 8-10 minutes. The HAADF–STEM, EDS mappings of samples on dried copper grids with the cross-sectional hybrids and dispersed Zn-doped-CDs were carried on transmission electron microscope (TEM, FEI Talos F200X, America) at 200 kV.

### Redox potential and Zeta potential analysis

The *Syne* cells were collected and resuspended in fresh BG11 medium to an OD_730_ of 9. As for CDs–*Syne* hybrid, 100 mg L^−1^ CDs was added to the *Syne* cells solution and incubated for 1 hour. The redox potentials of CDs (100 mg L^−1^), *Syne* cells and CDs– *Syne* hybrid were measured on a PHS-3C laboratory pH meter with a 501 OPR composite electrode (INESA Scientific InstrumentCo., Ltd, Shanghai, China).

CDs (100 mg L^−1^), *Syne* cells and CDs–*Syne* hybrid were diluted 200 times before measuring the Zeta potential. Zeta potentials of those diluted samples were analyzed on Zetasizer (Malvern ZSU3200, Britain).

### Photoelectrochemical analysis

Photoelectrochemical analysis was measured using electrochemical workstation (CHI1000C, Chenhua, Shanghai, China) through a standard three-electrode system in 10 mM phosphate buffered saline (PBS) electrolyte. A platinum wire and Ag/AgCl (3 M NaCl) served as the counter electrode and reference electrode, respectively. The working electrodes were prepared as following procedure: (1) Dropped 30 μL of 500 mg L^−1^ CDs onto a carbon paper electrode (1×1 cm^2^) and vacuum drying to form the CDs working electrode; (2) *Syne* cells were cultured in BG11 medium to an OD_730_ about 2. The *Syne* cells were collected and resuspended in fresh BG11 medium to an OD_730_ of 9. Dropped 30 μL of *Syne* cells solution to a carbon paper electrode (1×1 cm^2^) and vacuum drying to form the *Syne* working electrode; (3) The *Syne* cells solution (OD_730_ about 9) was prepared the same above. Added 500 mg L^−1^ CDs into the cell solution and incubated for 1 hour for constructing the CDs–*Syne* hybrid system. Dropped 30 μL of the hybrid system onto a carbon paper electrode (1×1 cm^2^) and vacuum drying to form the CDs–*Syne* hybrid electrode. The photocurrent measurements were performed using 50 mW cm^−2^ LED white light in 10 mM PBS electrolyte. The current density was recorded under 0 V bias vs Ag/AgCl with the light pulse (10 seconds on and 10 seconds off).

### Transient absorption experiment

The transient absorption set-up and experiment was performed according to our previous study^48^. The delay of probe was up to ∼2 ns and can be tuned by a delay line. The size of pump beam and probe are ∼590 µm and ∼200 µm, respectively. The samples are measured in the cuvette with 2mm optical pathlength. Pump light with an energy of 3.1 eV was used to excite CDs, and the electron dynamics signal in the CDs was observed in the 435-440 nm range of the probe light.

To quantitatively represent the transfer rate of charges, a biexponential function was used to fit the TA data:

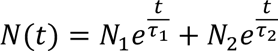

Where *N*(*t*) is the carrier population at the pump-probe delay of *t*, *τ*_1_ and *τ*_2_ represents the charge transfer process and charge recombination process. The charge transfer process times for samples were obtained by weighting *τ*_1_ and *τ*_2_ with *N*_1_and *N*_2_, respectively.

### The growth of *Syne* with CDs addition

*A. S. elongatus* PCC 2973 cells were inoculated with an initial OD_730_ of 0.3 in BG11 medium. CDs were added to the cultured medium with specifical concentration (0, 10, 40, 80 and 160 mg L^−1^, respectively). The shake flask cultures were performed at 38 ℃ under white light (5.5 mW cm^−2^) in 100 mL sealed flasks with 20 mL of BG-11 medium containing 100 mM NaHCO_3_. The growth of *S. elongatus* PCC 2973 was monitored by measuring the absorbance at 730 nm (OD_730_) at different time points. Specific growth rate was calculated using the equation :

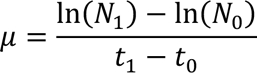

where t_1_ and t_0_ were cultivation time and N_1_ and N_0_ were the OD_730_ at t_1_ and t_0_, respectively.

### Oxygen evolution

The photosynthetic oxygen evolution rate of *Syne* and CDs–*Syne* were determined with a Clark-type oxygen electrode (Hansatech Chlorolab 2) according to previous reports^26^. *Syne* cells were collected from the exponential growth phase and resuspended in 2 mL BG11 medium containing 100 mM bicarbonate to an OD_730_ of 2. CDs were added to the culture medium solution with specifical concentration (0, 10, 40, 80 and 160 mg L^−1^, respectively). The oxygen evolution rate was firstly recorded under the light intensity of 30 mW cm^−2^ at 25 °C. To further explore the maximum oxygen evolution rate of *Syne*, the oxygen evolution rate was recorded with 40 mg L^−1^ CDs under different light intensity (7.5, 15, 30, 45, 60 mW cm^−2^).

### The chlorophyll fluorescence of photosystems

The chlorophyll fluorescence parameters were measured using a pulsed amplitude modulation (PAM) fluorimeter (Dual-PAM 100, Walz) as previously reported^26, 27^. *Syne* cells were collected from the exponential growth phase and resuspended in 2 mL BG11 medium containing 100 mM bicarbonate to an Chl a concentration at 20 mg mL^−1^. Cells were first incubated in dark conditions for 10 minutes. The light response curves were monitored to get the chlorophyll fluorescence kinetic parameters, including the relative electron transport rate (rETR (II) and rETR (I)) and the effective quantum yield (ϕPSII and Y(I)).

### NADP^+^ and NADPH measurements

The intracellular concentration of NADP^+^ and NADPH were measured using NADP^+^/NADPH Assay Kit (Beyotime, Cat. No. S0179). 40 mL *Syne* cells (OD_730_=0.3) with CDs (40 mg L^−1^) or without CDs were firstly inoculated for 1 hour in dark, followed by light (5.5 mW cm^−2^) for 15 mins. The cell pellets were collected, washed twice with cold PBS, and resuspended in cold PBS. The solution was divided into two vials (with the same volume). The precipitate was collected and resuspended with 0.5 ml of 0.2 M HCl (for NADP^+^ extradition) or 0.5 ml of 0.2 M NaOH (for NADPH extraction). The samples were first bathed in water at 55 °C for 10 min, then cooled on ice for 5 min, then neutralized with 0.5 ml of 0.1 M NaOH or HCl. Samples were then vortexed at high speed. After incubation on ice for 10 min, the samples were centrifuged at 10,000 rpm for 5 min and the supernatants were then used for assay. The concentration of NADP^+^ and NADPH in the solution was measured by the NADP+/NADPH Assay Kit. The intracellular concentrations of NADP^+^ and NADPH were calculated based on the used biomass.

### Metabolite measurements

*Syne* cells with or without CDs were grown in shake flasks with 100 mL BG-11 medium containing 100 mM NaHCO_3_ under white light (5.5 mW cm^−2^) to OD_730_ of ∼1. Cells were collected by fast filtration on the filters. Metabolism was quenched, and metabolites were extracted by rapid transfer of the filters into –20 ℃, 40:40:20 % acetonitrile/methanol/ water with 0.1% formic acid. After incubation at – 20 °C for 20 minutes, the samples were centrifuged, and the supernatant was collected. For ATP measurement, samples were collected by centrifugation at 8000 rpm for 2 min after 15 min of light exposure, followed by rapid quenching and extraction.

The metabolites extracted from the cells were analyzed by ultrahigh performance liquid chromatograph (UltiMate 3000, Thermo Fisher) coupled to a quadrupole-orbitrap mass spectrometer (Q-Exactive, Thermo Fisher). The injection volume was 10 μL. Solvent A was 20 mM ammonium acetate adjusted to pH 9.5 with ammonium hydroxide, and solvent B was acetonitrile. Metabolites were separated with a Luna NH_2_ column (100 mm × 2 mm, 3 μm particle size, Phenomenex). The column was maintained at 15℃ with a solvent flow rate of 0.3 mL min^−1^, and the gradient of B was as follows: 0 min, 85%; 10 min, 45%; 15 min, 2%; 18 min, 2%; 18.1 min, 85%; 24 min 85% B. The mass spectrometer with a heated electrospray ionization source was operated in positive and negative modes. The key parameters were as follows: ionization voltage, +3.8 kV/–3.0 kV; sheath gas pressure, 35 arbitrary units; auxiliary gas, 10 arbitrary units; auxiliary gas heater temperature, 350 °Capillary temperature, 320 °C. The mass spectrometer was run in full scan mode at an m/z 70–1,000 scan range and 70,000 resolutions. Data processing and ion annotation based on accurate mass were performed in Xcalibur 4.0 (Thermo Fisher) and Compound Discoverer 2.0 (Thermo Fisher). A subset of identified compounds was verified by mass and retention-time match to authenticated standards.

### Isotope labeling experiment

Isotope labeling experiments were performed with ^13^C-labeled NaHCO_3_ ([^13^C]-bicarbonate) as tracers. Labeled compounds were ≥99% pure and purchased from Cambridge Isotope Lab. *Syne* cells with or without CDs (40 mg L^−1^) were grown in shake flasks with 100 mL BG-11 medium containing 100mM NaHCO_3_ under continuous light (5.5 mW cm^−2^) to OD_730_ of ∼1. Cells were collected by fast filtration and transferred to BG-11 plates containing 24 mM [^13^C]-bicarbonate for isotope labeling experiment. At various time points after the isotope addition, Metabolism were fast quenched for LC–MS analysis of labeling of metabolites.

### CO_2_-fixation-pathway fluxes

To quantify the CO_2_-fixation fluxes through Rubisco in the CB cycle, cells were switched to medium containing [^13^C]-bicarbonate, and the dynamic labeling patterns of PEP, RuBP, and 3-phosphoglyceric acid (PGA) were measured as described above using LC-MS. Mass isotopomer distributions (MIDs) of metabolites were calculated from measured peak areas of the mass spectra and corrected for naturally occurring ^13^C. Model construction depended on the mapping of carbon atoms between substrates and products of the biochemical reactions composing the metabolic node, which is described in the following sections. The model was then used to identify the set of parameters including the fluxes or flux ratio of interest, which was compatible with the measured labeling patterns of metabolites. Parameters were fitted by minimizing the weighted sum of squared residuals (*χ*2) between measured and simulated labeling kinetics^56^. All calculations were performed in Matlab 7.8.0 (Mathworks). The flux through Rubisco was estimated with following equations:

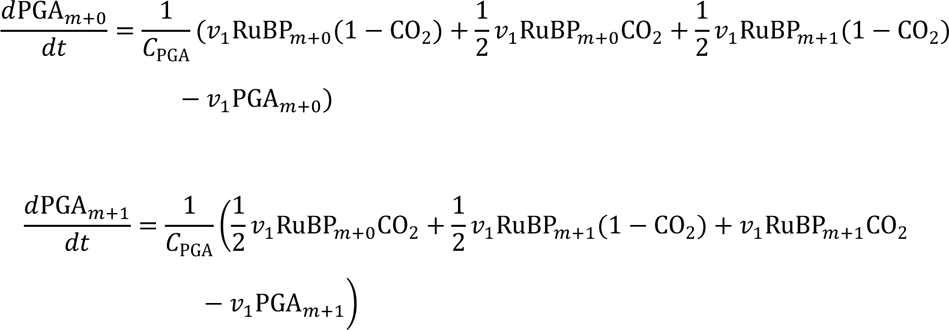

where *v*_1_ is flux from RuBP to PGA, C_PGA_ is PGA concentration and CO_2_ is ^13^C labeling degree of CO_2_.

### Glycerol fermentation of engineering *S. elongatus* 7942 XG608

The glycerol producing strain XG608^48^ was cultured at 30℃ under continuous LED white light (1.35 mW cm^−2^) in BG11 medium containing 100 mM NaHCO_3_ and 20 mM HEPES. IPTG (1 mM) was added to the cultures at OD_730_ of 0.4. After cells were grown to the exponential phase (OD_730_ about 1.0-1.5), the *Syne* cells were collected, washed, and resuspended in fresh BG11 medium to an OD_730_ of required value. Added 100 mg L^−1^ CDs into the *Syne* cultures and incubated 30 mins under 1.35 mW cm^−2^ white light. Added same volume of solvent (ethanol) into the *Syne* cultures as the control group. The glycerol producing experiments were performed as following: (1) with the required OD_730_ under the same LED light intensity of 1.35 mW cm^−2^ in Figure 3j and 3k; (2) under different LED light intensity with the same OD_730_ of 1 in Supplementary Figure 4; (3) under simulated solar light with full-spectrum and visible light only (using the filter) at the intensity of 1.35 mW cm^−2^ by the same OD_730_ of 1 in Figure 4c and 4d. For the inhibitor’s experiments, the inhibitor (10 µM DCMU, 10 µM DBMIB, 200 µM PMA, 10 µM trifluoromethoxy carbonylcyanide phenylhydrazone (FCCP), and 5 mg L^−1^ Antimycin A) was added together with CDs materials, respectively. The incubation and glycerol production were performed as the same above.

### Glycerol measurements

The samples were centrifuged for 10 minutes at 15000 rpm and the supernatants were used for glycerol measurements by glycerol assay reagent (APPLYGEN, E1002-250, China). Briefly, the 25 µL supernatant was added to 75 µL working solution and incubate 30 mins at 30 ℃. The glycerol productions were analyzed by measured the absorption of the reaction solution at the wavelength of 550 nm.

### The growth conditions of *A. thaliana*

*A. thaliana* seeds (Col-0 ecotype) were sterilized with 75% ethanol for 10 minutes. After rinsing with water and drying at room temperature, the seeds were placed on 1/2 MS medium (PhytoTech, M524). The seeds were kept in the dark at 4 °C for 3 days and then placed at 22 °C for germination. One week later, the seedlings were transplanted into the seedling block under the growth conditions (temperature: 22 °C; relative humidity 50-70%; light intensity: 120 μmol m^−2^ s^−1^, 12 hours light /12 hours darkness) for another two weeks. The seedlings with consistent growth were chosen for CDs treatment.

CDs was dissolved in a small amount of ethanol and then diluted to 10 mg L^−1^, 50 mg L^−1^, and 100 mg L^−1^, respectively. CDs was sprayed on the surface of the *A. thaliana* (10 samples for each group) leaves at the concentrations of 0 mg L^−1^ (as control), 10 mg L^−1^, 50 mg L^−1^, and 100 mg L^−1^. The CDs treatment was repeated every two days (seven times in total). All the plants were placed in the dark for 1 hour after CDs spraying.

To measure the fresh weight and area of leaves in *A. thaliana*, the leaves were clipped and any soil on them was wiped off quickly. The fresh weight of leaves was weighed by an analytical balance (Mettler Toledo, ME802E). The area of the leaves was measured using ImageJ software. The significant differences between samples and the corresponding controls were analyzed using two-tailed Student’s *t*-test for pairwise comparisons, one-way ANOVA analysis with Tukey’s multiple comparison test as specified in the Figure legends. Samples sharing lowercase letters are not significantly different.

## Data availability

All data presented in this manuscript are available in the paper and its Supplementary Information.

## Acknowledgements

We acknowledge the Shenzhen Infrastructure for Synthetic Biology for instrument support and technical assistance. This work was supported by the National Key R&D Program of China (Grant No. 2021YFA0909700), the National Natural Science Foundation of China (Grant No. 32230060, 32171426, 31925001, 32070564, and 22373046), Shenzhen Science and Technology Program (Grant No. JCYJ20220818101804010, RCYX20221008092901004, TCYJ20220531100006011, and JCYJ20220818100212027), the Chinese Academy of Sciences (Grant No. XDB27020000), Yunnan Fundamental Research Projects (Grant No. 202301BF070001-013 and 202201AT070090), and Xingdian Talent Support Program of Yunnan Province (Grant No. C619300A094).

## Author Contribution

The project was conceptualized by X.G. and C. Y., and was supervised C. Y., X. G., J. L., Y. L., and X. C.. X. W., J. D., H. H. prepared the CDs and performed the CDs, CDs– *Syne* hybrid characterizations. W. C. performed the experiments of *Syne* growth, oxygen evolution, photosystem activity, CO_2_ fixation flux, and intracellular metabolites measurements. X. Y. performed the transient absorption experiment. H. H. performed the glycerol production experiments. Y. Y. performed the experiments of *Arabidopsis thaliana*. X. G., C. Y., Y. L., X. C., X. W., W. C., H. H., and Y. Y. wrote the manuscript with input from all authors. W. Z., Q. Z., and R. L. A. analyzed the data and edited the manuscript. All authors discussed the results and commented on the manuscript.

