## Supplementary Information 1 for "Enhancing Plant Photosynthesis using Carbon Dots as Light Converter and Photosensitizer"

### Supplementary Figures

**Supplementary Figure 1** The synthesis and characterizations of CDs.

**Supplementary Figure 2** The characterizations of CDs–*Syne*.

**Supplementary Figure 3** The transient absorption spectroscopy analysis of CDs

**Supplementary Figure 4** The intracellular metabolite concentration in *Syne* and CDs–*Syne*.

**Supplementary Figure 5** The effect of light intensity on glycerol production in CDs–*Syne*.

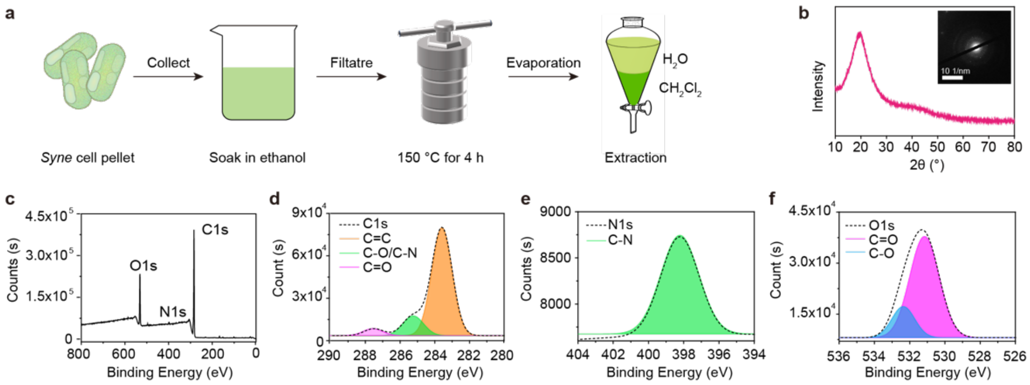


**Supplementary Figure 1: The synthesis and characterizations of CDs. a.** Schematic of CDs synthesis by hydrothermal method from *Syne* cell pellet. **b.** The XRD spectrum of CDs, and illustration is the TEM diffraction patterns, indicate CDs have the amorphous structure. **c-f.** XPS survey spectra of CDs. Deconvolution spectra of d. C1s, e. N1s and f. O1s for CDs.

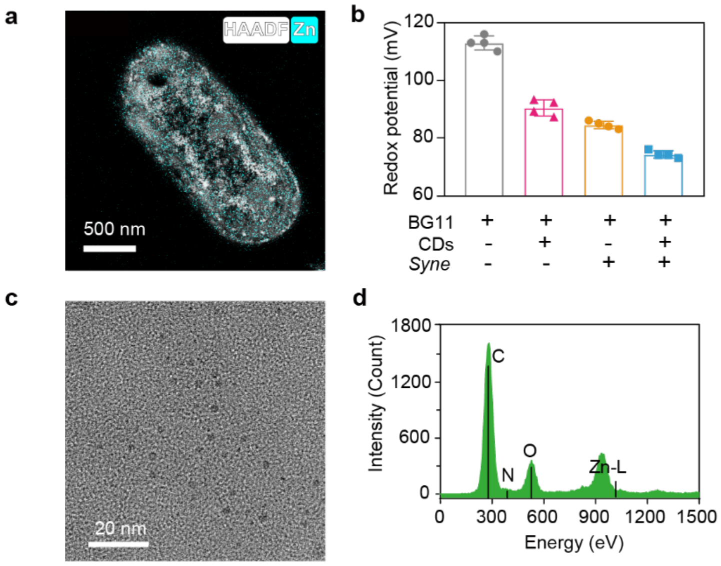


**Supplementary Figure 2: The characterizations of CDs–*Syne*. a.** The distribution of CDs in CDs–*Syne* hybrid system. High-Angle Annular Dark Field (HAADF)-STEM and energy dispersive x-ray spectroscopy (EDS) mapping showing that CDs were mainly distributed in the cells. CDs doped with Zn was used to prepare hybrid system, and EDS imaging was based on Zn element. **b.** The redox potential of BG11 medium, CDs, *Syne* and CDs–*Syne*. The potential of CDs–*Syne* is higher than that of Syne or CDs, indicating the formation of hybrid system. **c.** TEM morphology of CDs doped with Zn. **d.** EDS mapping of CDs doped with Zn.

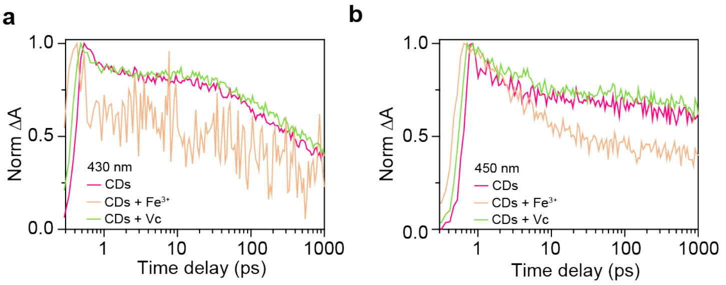


**Supplementary Figure 3: The transient absorption spectroscopy analysis of CDs. a-b.** TA spectra of CDs with $Fe^{3+}$ or vitamin c (Vc). Vc serves as a hole trapping agent, while $Fe^{3+}$ is used for electron capture. The similar spectra of CDs and CDs with Vc suggested that the photocarriers of CDs are mainly electrons, with a relatively small proportion of holes.

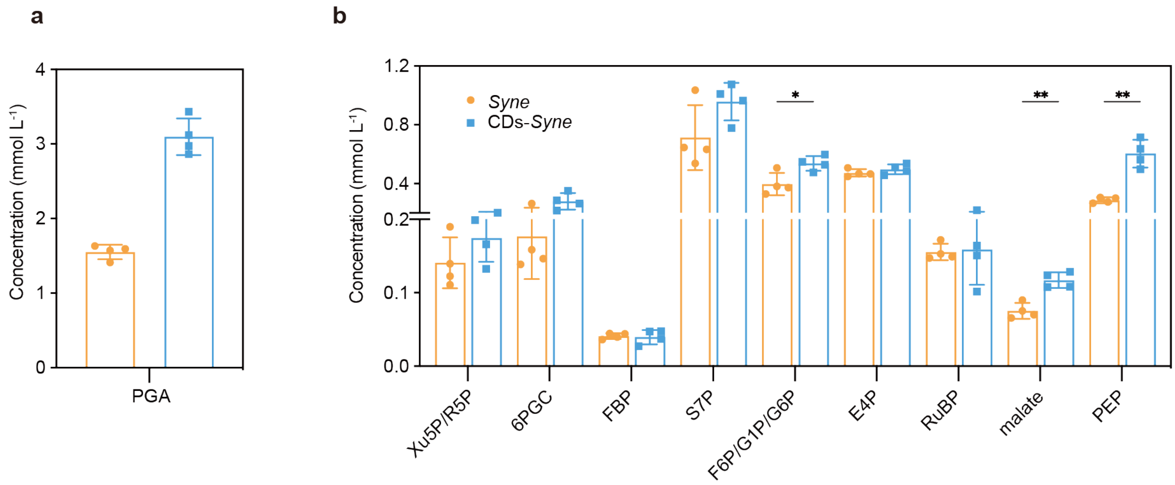


**Supplementary Figure 4: The intracellular metabolites concentration in *Syne* and CDs–*Syne*. a** and **b.** Concentration of intracellular metabolites of central carbon metabolism in *Syne* and CDs–*Syne*. PGA, 3-phosphoglyceric acid; Xu5P, xylulose 5-phosphate; R5P, ribose 5-phosphate; 6PGC, 6-phospho-gluconate; FBP, fructose 1,6-bisphosphate; S7P, 7-phosphosedoheptose; F6P, fructose 6-phosphate; G1P, glucose 1-phosphate; G6P, glucose 6-phosphate; E4P, erythrose 4-phosphate; RuBP, ribulose-1,5-bisphosphate; PEP, phosphoenolpyruvate.

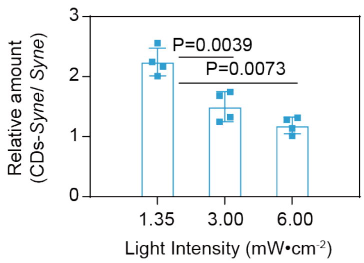


**Supplementary Figure 5: The effect of light intensity on glycerol production in CDs–*Syne*.** Relative glycerol production by CDs–*Syne* hybrids compared to that by *Syne* cells under different light intensities (1.35, 3, and 6 mW $cm^{-2}$).
